# Competitive binding of MatP and topoisomerase IV to the MukB dimerization hinge

**DOI:** 10.1101/2021.03.03.433707

**Authors:** Gemma L. M. Fisher, Jani R. Bolla, Karthik V. Rajasekar, Jarno Mäkelä, Rachel Baker, Man Zhou, Josh P. Prince, Mathew Stracy, Carol V. Robinson, Lidia K. Arciszewska, David J. Sherratt

## Abstract

SMC complexes have ubiquitous roles in chromosome organisation. In *Escherichia coli,* the interplay between the SMC complex, MukBEF, and matS-bound MatP in the replication termination region, ter, results in depletion of MukBEF from ter, thus promoting chromosome individualisation by directing replichores to separate cell halves. MukBEF also interacts with topoisomerase IV ParC_2_E_2_ heterotetramers, to direct its chromosomal distribution to mirror that of MukBEF, thereby facilitating coordination between chromosome organisation and decatenation by topoisomerase IV. Here we demonstrate that the MukB dimerization hinge binds ParC and MatP with the same dimer to dimer stoichiometry. MatP and ParC have an overlapping binding interface on the MukB hinge, leading to their mutually exclusive binding. Furthermore, the MukB hinge fails to stably associate with *matS*-bound MatP, while MatP mutants deficient in *matS* binding are impaired in MukB hinge binding, demonstrating that *mats* competes with the hinge for MatP binding. Cells expressing MukBEF complexes containing a mutation in the MukB hinge interface for ParC/MatP binding are deficient in ParC binding *in vivo,* despite having a Muk^+^ topoisomerase IV^+^ phenotype. This mutant protein is also impaired in MatP binding *in vitro,* and cells expressing this variant exhibit a MukBEF cellular localisation consistent with impaired MatP binding.

## INTRODUCTION

In *Escherichia coli* and its *γ*-proteobacterial relatives, the Structural Maintenance of Chromosomes (SMC) complex, MukBEF, interacts with the decatenase topoisomerase IV (topoIV) and MatP, the 800 kb replication terminus region (ter) binding protein, to organise the chromosome and to ensure that newly replicated sister chromosomes are decatenated and individualised, so that they can be segregated to opposite cell halves (1–3). The details of how these processes act mechanistically and how they are coordinated remain to be determined.

Although MukBEF architecture exhibits many of the conserved features present in prokaryote and eukaryote SMC complexes (Figure 1A), its dimeric kleisin, MukF, can direct the formation of dimer of dimer complexes that have been inferred to act *in vivo* (4) and form *in vitro* (5). It has been proposed that *in vivo,* MukBEF action forms, likely through loop extrusion, a MukBEF axial core, from which DNA loops of 15-50 kb emanate (2). In MatP^+^ cells, MukBEF is depleted from the ~800 kb *ter* region, subject to MatP binding to 23 *matS* (13 bp) sites distributed throughout *ter* (2, 3, 6). In the absence of MatP, MukBEF forms axial cores around the whole chromosome, leading to chromosome rotation, mis-regulation of decatenation, and aberrations in the patterns of chromosome segregation (2, 3). *The E. coli* MukB dimerization hinge interacts specifically with the ParC subunit of topoIV, stimulating its catalytic activity and presumably directing it to sites of MukBEF action on the chromosome (7–12). Consistent with this, the cellular copy numbers of MukBEF and topoIV are comparable (11). Furthermore, an initial characterisation showed that MatP also binds the MukB dimerization hinge (6).

**Figure 1.**
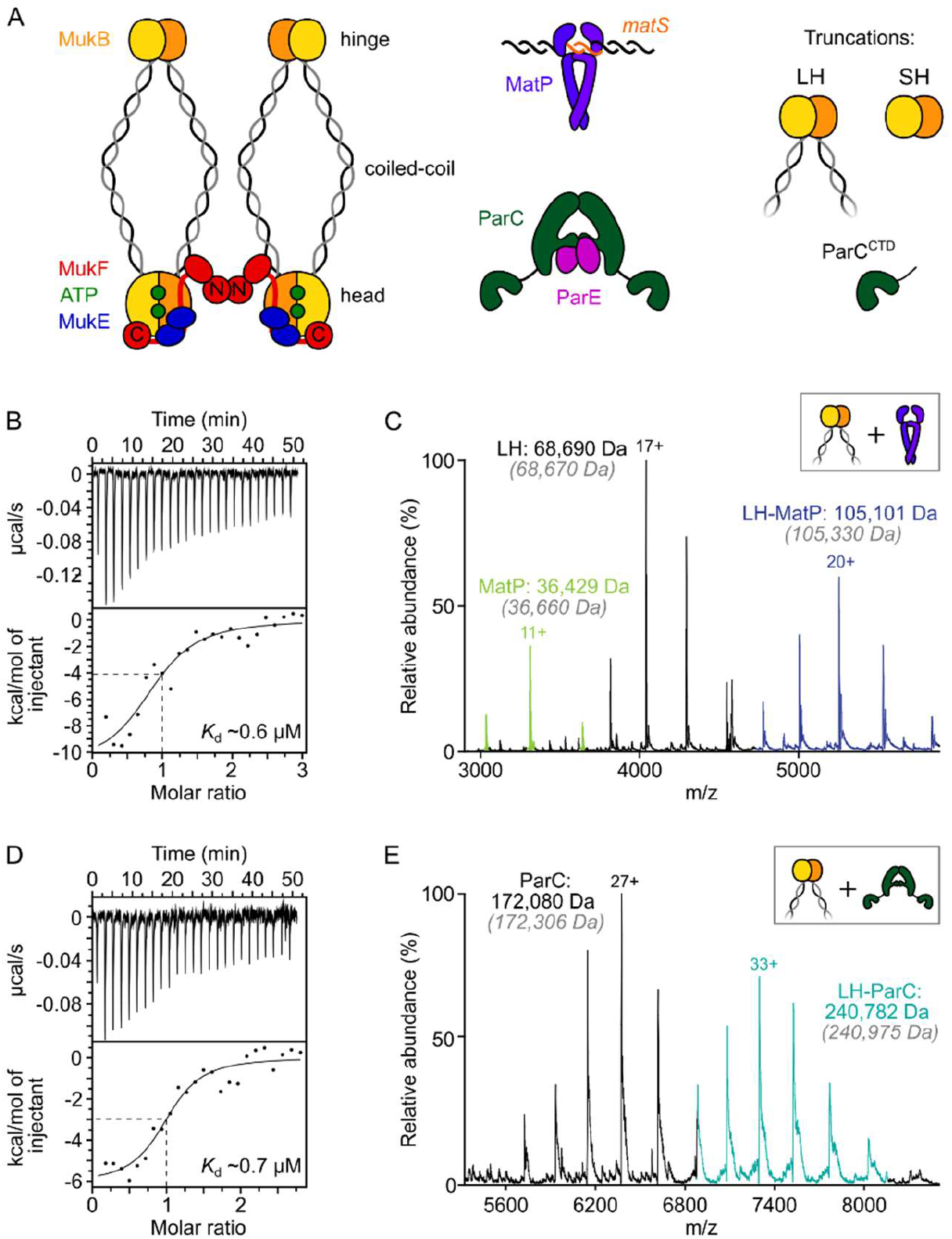
ParC dimers and MatP dimers each bind the dimeric MukB hinge with a 1:1 stoichiometry. (**A**) Schematics of the functional unit of MukBEF complexes, the topoIV heterotetramer and MatP-matS complex. (**B**) ITC raw thermogram and binding isotherm following titration of 100 μM MatP into 10 μM LH at 25 °C. Fitted parameters for a one-site model from three independent measurements were n = 0.96 ± 0.08, *K*_d_ = 0.59 ± 0.04 μM, AH = −16.8 ± 2.2 kcal/mol and AS = −27.8 cal/mol/deg. Reported errors are fitting errors. (**C**) nMS. MatP was incubated at a 2-fold molar excess with LH before injection. An example spectrum is shown with the only detected LH-MatP complex highlighted in purple. Grey italics denote the theoretical mass of complexes. (**D**) ITC raw thermogram and binding isotherm following titration of 100 μM LH into 10 μM ParC at 25 °C. Fitted parameters for a one-site model from three independent measurements were n = 0.84 ±0.11, *K*_d_ = 0.69 ± 0.11 μM, AH = −7.48 ± 1.3 kcal/mol and AS = −3.12 cal/mol/deg. Reported errors are fitting errors. (**E**) nMS. ParC was incubated at a 2-molar excess with LH before injection. An example spectrum is shown with the only LH-ParC complex (1:1) detected highlighted in cyan.

Here, we undertook an extensive characterization of the binding of MatP and topoIV to the MukB hinge *in vitro.* Using a range of independent ensemble and single-molecule assays, we showed that a dimeric MukB hinge binds a MatP dimer, a ParC dimer, or a single topoIV heterotetramer (ParC2E2) with comparable affinities. Importantly, we demonstrated that binding of MatP and ParC (or topoIV) to the MukB hinge is mutually exclusive, with them interacting with an overlapping interface on the hinge, thereby supporting an earlier inference of competitive binding from *in vivo* analyses (6, 12). Cells expressing the MukB variant, which no longer interacts with topoIV and MatP *in vitro,* had a Muk^+^ topoisomerase IV^+^ phenotype *in vivo.* Quantitative live cell imaging showed that the variant hinge no longer interacts with ParC *in vivo* and such cells had a phenotype consistent with them also being impaired in the interaction with MatP. Surprisingly, interaction of MatP with the MukB hinge *in vitro* was inhibited in the presence of matS-containing DNA; given that the displacement of MukBEF complexes from *ter in vivo* requires that the MukBEF complexes interact, at least transiently, with MatP-matS. Consistent with the observation that *matS* binding to MatP inhibits its interaction with the MukB hinge, mutants in MatP that fail to bind *mats,* were impaired in hinge binding.

## MATERIALS AND METHODS

### Protein overexpression and purification

Two MukB hinge-based constructs were used: long hinge (LH, 568-863) and short hinge (SH, 667-779). SH was purified using a C-terminal 6xHis-tag whereas LH was initially purified as an N-terminal fusion to MBP.

SH-His was overexpressed and first purified by TALON affinity chromatography as described for MukB (13). Then, peak fractions were diluted to 100 mM NaCl and loaded onto a 5 ml HiTrap Q XL column (GE Healthcare) equilibrated in 50 mM HEPES pH 7.3, 100 mM NaCl, 1 mM EDTA and 10% (v/v) glycerol. Elution was achieved over 40 column volumes using a gradient of 100-1000 mM NaCl. Appropriate fractions were pooled and concentrated by centrifugal filtration (Vivaspin 20, 5,000 MWCO PES, Sartorius) for loading onto a Superose 6 Increase 10/300 GL (GE Healthcare) column equilibrated in storage buffer (50 mM HEPES pH 7.3, 300 mM NaCl, 1 mM EDTA, 1 mM dithiothreitol (DTT) and 10% (v/v) glycerol). Peak fractions were assessed for purity (>90%) by sodium dodecyl sulphate-polyacrylamide gel electrophoresis (SDS-PAGE; 4-20% gradient) and Coomassie staining, concentrated as appropriate by centrifugal filtration, and snap frozen as aliquots for storage at −80 °C.

MBP-LH (and derivatives thereof) was overexpressed using the pMAL-c5X vector system according to the vendor-supplied protocol in NEBExpress cells (New England Biolabs). Glucose was present at 0.2% (w/v) throughout overexpression to repress amylase expression. For purification, cells were resuspended in lysis buffer (50 mM HEPES pH 7.3, 300 mM NaCl, 5% (v/v) glycerol) supplemented with a protease inhibitor cocktail (EDTA-free, Thermo Scientific Pierce) and mechanically lysed using a homogeniser. The lysate was clarified by centrifugation and the cleared suspension diluted 4-fold in cold lysis buffer per 25 ml of extract. This was added to 5 ml (per 2 L of culture) amylose resin (New England Biolabs) equilibrated in lysis buffer and left on a tube roller shaker at 4 °C for 1 h. This suspension was loaded onto a gravity-flow column. The settled resin was washed with 10 column volumes of lysis buffer. MBP-LH was eluted in 2 column volumes using lysis buffer supplemented with 10 mM maltose. MBP was cleaved from LH using Factor Xa (New England Biolabs) at a w/w ratio of 1% Factor Xa:LH. Efficient cleavage typically required incubation at 4 °C for 36 h. The cleaved sample was diluted until a final [NaCl] of 100 mM and then loaded onto a 5 ml HiTrap Q XL column equilibrated in 50 mM HEPES pH 7.3, 1 mM EDTA, 10% (v/v) glycerol with 100 mM NaCl. Elution was achieved with a linear 40 column volume gradient to 1 M NaCl to isolate tagless LH and remove Factor Xa. Finally, appropriate fractions were pooled for further purification and buffer exchange by SEC as described for aforementioned for MukB.

Purification of full-length ParC and MatP (MatPΔ18C;1-132) used an N-terminal and C-terminal 6xHis-tag, respectively. Overexpression and initial purification was completed as previously published (6) with the addition of a SEC step as described above. Note, these proteins are unstable during purification when using buffers with a NaCl concentration below 300 mM. ParC R705E R729A and MatP K71E/A, Q72E/A, R75E/A, R77E/A variants were produced by site-directed mutagenesis (Q5 Site-Directed Mutagenesis Kit, New England Biolabs), verified by sequencing, and purified using the same protocol.

ParC^CTD^ used an N-terminal MBP fusion for ease of purification and to improve stability of the construct as reported by Vos *et al.* (10, 14). MBP-ParC^CTD^ was overexpressed and initially purified using amylose affinity chromatography as described for MBP-LH, however the MBP fusion was not removed. Instead, MBP-ParC^CTD^ was loaded onto a Superose 6 Increase 10/300 GL column (GE Healthcare) equilibrated in storage buffer.

ParE-His was overexpressed and initially purified using TALON affinity chromatography as for MukB (13). The eluate was diluted to 100 mM NaCl and loaded onto a 1 ml HiTrap Q XL column equilibrated in 50 mM HEPES pH 7.3, 100 mM NaCl, 1 mM EDTA and 10% (v/v) glycerol. Elution was achieved over a 20-column volume gradient to 1 M NaCl. Selected fractions were passed over a Superose 6 Increase 10/300 GL (GE Healthcare) column equilibrated in storage buffer. TopoIV reconstitution required mixing of equimolar amounts of ParC and ParE which were incubated for 30 min on ice. Efficient reconstitution under these conditions was verified in analytical SEC.

### Fluorophore labelling of MatP and MukB

Endogenous MatP contains no native cysteines; a single cysteine was engineered into the linker between MatP and its 6xHis-tag. Immediately following purification, MatP-Cys-His was treated with 0.2 mM TCEP for 30 min at 22 ± 1 °C. Cy3B maleimide was dissolved in anhydrous DMSO to produce a 10 mM stock and immediately added to MatP-Cys-His at a 6-fold molar excess. This reaction was rotated end-over-end at 4 °C for 16 h whilst protected from light and then quenched by addition of DTT to a final concentration of 5 mM. Excess dye was removed by SEC. Labelling efficiency was calculated (89%) by spectrophotometry and using a vendor-supplied correction factor of 0.08 for Cy3B absorbance at 280 nm.

Unnatural amino acid labelling was used for conjugating fluorophores to MukB. S718 was mutated to an amber stop codon in a pBAD24 expression vector. This was co-transformed with pEVOL-pAzF (Addgene) into an *E. coli* C321.ΔAeluate was incubated with 125 strain (15, 16), where endogenous MukB has a C-terminal 3xFLAG tag, and UAG has been reassigned as a sense codon (FW01). 1% (w/v) glucose was present throughout overexpression which was induced by the addition of L-arabinose to 0.4 % (w/v) and p-azidophenylalanine (azF) to 1 mM at an OD_600_ of 0.6. Expression proceeded for 4 h at 30 °C. azF-MukB was purified as for the wild type protein (13) with an additional step post-TALON resin, where the eluate was incubated with 125 μl of equilibrated ANTI-FLAG M2 agarose affinity gel (Sigma) for 1 h on a rolling shaker at 4 °C before being poured onto a column. The flow-through was recovered and processed as for wild type protein. A 20-fold excess of the dye (DBCO-TAMRA) was added to MukB at ~15 uM. The reaction was left to proceed for 1 h at 22 ± 1 °C and then moved to 4 °C for 16 h in the absence of light. Free dye was removed from labelled MukB by SEC using a Superdex 200 10/300 GL column (GE Healthcare). Labelling efficiency (typically 40-70%) was calculated using the molar extinction coefficient of the TAMRA dye at 547 nm (92,000 M^-1^cm^-1^), and a 0.3 correction factor for absorption at 280 nm by the dye. The suitability of substitution of S718 to a phenylalanine analogue was verified *in vivo* by assaying its ability (as S718F) to rescue the temperature-sensitive growth defect of a *ΔmukB* strain as previously described (13), and *in vitro* by measuring the ATPase activity of azF-MukB.

### DNA preparation

For native mass spectrometry, 50 bp matS-containing or non-specific double-stranded DNA of the same size and GC content was prepared by slowly annealing complementary oligonucleotides: 50 bp *matS* oligo 1 5’ CAG AGT TAA TCA GAA CGG TGA CAA TGT CAC AAA GAA AAA GAA CCT GTG CG 3’; 50 bp *matS* oligo 2 5’ CGC ACA GGT TCT TTT TCT TTG TGA CAT TGT CAC CGT TCT GAT TAA CTC TG 3’; 50 bp non-specific oligo 1 5’ CAG AGT TAA TCA CAA CGG TTC TCG ATC ATC AAA GAA AAA CAA GCT GTG CG 3’ and 50 bp non-specific oligo 2 5’ CGC ACA GCT TGT TTTT CTT TGA TGA TCG AGA ACC GTT GTG ATT AAC TCT G 3’. All oligos were dissolved in annealing buffer (10 mM Tris-HCl pH 8.0, 10 mM NaCl and 1 mM EDTA), mixed at an equimolar ratio, heated at 95 °C for 5 min and cooled slowly in 0.1 °C increments to 10 °C over 6 h. Double-stranded DNA formation was assessed by agarose gel electrophoresis.

For fluorescence correlation spectroscopy, 15 bp matS-containing or non-specific DNA hairpins were produced by resuspending the following oligonucleotides in annealing buffer to produce 50 μM stocks: 5’ GTG ACA ATG TCA C TTC CCT G TGA CAT TGT CAC 3’ and 5’ GTT CTC GAT CAT C TTC CCT G ATG ATC GAG AAC 3’ for *matS* and non-specific DNA, respectively. Both DNAs were heated at 95 °C for 15 min and then immediately placed in an ice-water bath for 10 min. Selective hairpin formation was assessed by PAGE using 15% TBE (pH 7.4) gels later stained with 0.5 μg/ml ethidium bromide 1X TBE solution.

### Circular Dichroism (CD) spectroscopy

CD spectra were collected at 20 °C using a 0.1 cm quartz cuvette in a JASCO J-815 spectropolarimeter equipped with a JASCO CDF-426S Peltier temperature controller. 0.05-0.1 mg/ml samples were buffer exchanged into 10 mM phosphate buffer pH 8.0 and 20 mM NaCl. Data was acquired across a 190-250 nm absorbance scan using a band width of 1 nm, a data pitch of 0.1 nm, and scan rate of 100 nm/min. 9 scans were accumulated and averaged, and the data normalised to molar ellipticity by calculation of the cell path length and concentration of peptide bonds. A buffer only baseline was subtracted from all datasets.

### Isothermal Titration Calorimetry (ITC)

Binding was assayed in a Malvern PEAQ ITC instrument at 25 °C. Averages and standard deviations of the obtained parameters were obtained from triplicate experiments. Data were analysed using the manufacturer’s software assuming a single binding site model.

### Native Mass Spectrometry (nMS)

Prior to nMS analysis, individual proteins were buffer exchanged into 200 mM ammonium acetate pH 8.0 either by SEC or using Biospin-6 (BioRad) columns and introduced directly into the mass spectrometer using gold-coated capillary needles (prepared in-house). To reconstitute the complexes, buffer exchanged proteins were mixed in different ratios and incubated on ice for 10 min. Data were collected on a Q-Exactive UHMR mass spectrometer (ThermoFisher). The instrument parameters were as follows: capillary voltage 1.1 kV, quadrupole selection from 1,000 to 20,000 m/z range, S-lens RF 100%, collisional activation in the HCD cell 50-200 V, trapping gas pressure setting kept at 7.5, temperature 100-200 °C, resolution of the instrument 12500. The noise level was set at 3 rather than the default value of 4.64. No in-source dissociation was applied. Data were analysed using Xcalibur 4.2 (Thermo Scientific) and UniDec (17). The theoretical and measured masses of all constructs used in nMS experiments in this study are listed in Supplementary Table 1. Data collection for all spectra was repeated at least 3 times.

### Analytical Ultracentrifugation (AUC)

All analytical ultracentrifugation data were obtained on a Beckman XL-I using absorbance optics. MatP and LH were taken at 100 μM in 50 mM HEPES pH 7.5, 100 mM NaCl and 1 mM MgCl_2_. Sedimentation velocity experiments were carried out at 40,000 rpm using an AnTi60 rotor at 20 °C. Cells were scanned every 10 min at 280 nm. All data were analysed using SEDFIT (18).

### Fluorescence Correlation Spectroscopy (FCS)

FCS experiments were performed using a bespoke confocal microscope with continuous excitation at 532 nm (50 μW, Samba, Cobolt). Time traces were acquired for 30 s using a SPQR14 avalanche photodiode (PerkinElmer), and autocorrelation functions were produced in real-time using a Flex02-02D correlation card (Correlator.com). Data acquisitions were performed with custom software written in LabVIEW (National Instruments). Fluorescence arrival times were recorded on a SPQR-14 detector (PerkinElmer) and processed using custom software written in LabVIEW, MATLAB (MathWorks), and Python (Python Software Foundation).

Samples in the μM range (for fluorophore-labelled species) were deposited onto PEGylated slides in FCS buffer (20 mM Tris-HCl pH 7.5, 50 mM NaCl, 0.3 mg/ml BSA, 2 mM DTT, 0.05% (v/v) Tween-20). All buffers were UV-bleached before use. TMR-labelled MukB (conjugated at S718TAG using UAA as described above) was used to calculate the diffusion time of MukB alone. Competitor proteins were added at 2.5-50X molar excess. Complex samples with more than one binding component were incubated for 10 min at 22 ± 1 °C prior to data acquisition. Data for each sample was collected from >20 datasets.

For the single diffusing species, the auto-correlation function *G*(*τ*) is given by:

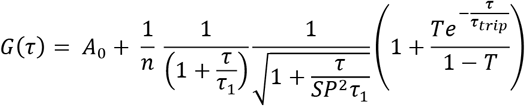

Where *A*_0_ is the offset, *n* is the effective number of particles in the confocal volume, *SP* is the structural parameter which describes elongation of the confocal volume, *T* is the fraction of MatP in triplet state. *τ*_1_ is the characteristic diffusion time of free MatP, and *τ_trlp_* is the characteristic residence time in triplet state.

For a two-component system, such as that consisting of free MatP and MukB-bound MatP, the correlation function becomes:

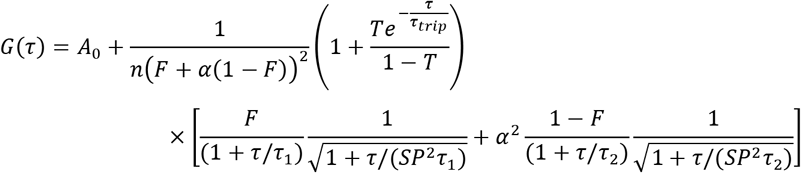

Where *A*_0_, *n, SP, T* and *τ_trlp_* are the same parameters as described above. *τ*_1_ is the characteristic diffusion time of free MatP, and *τ*_2_ is the characteristic diffusion time of bound-MatP (MukB-MatP). *F* is the fraction of molecules of MatP and *α* is the relative molecular brightness of MatP and MukB-MatP (regarded as 1).

FCS data were fitted using PyCorrFit software (19). A data range of 300-750 was used for channels setting which defined the timescale as 1.8000*10’^2^ to 3.0802*10^+2^ ms. The following constraints were set for fitting: *τ_trlp_* at 100-1000 ps and *τ*_1_ at 0-100 ms. For the single diffusing species fitting (MatP-Cy3B), *SP* was defined as the fixed parameter at default value of 5. For the two-component system fitting, *τ*_1_ was set as 4.5 ms for free MatP-Cy3B (diffusion time of ~ 4.5 ms). *SP* was the fixed parameter with a default value of 5, *τ_trlp_* was 100 – 1000 μ and *τ*_2_ as 20-100 ms. Binding data were fitted using the Hill equation with Origin software (version 2017, OriginLab Corporation).

### Native Polyacrylamide Gel Electrophoresis (PAGE)

Samples were prepared in 50 mM HEPES pH 7.5, 100 mM NaCl, 1 mM MgCl_2_, 1 mM DTT and 10% glycerol (v/v) buffer. In experiments used to monitor LH/LH^KKK^ binding to MatP and ParC, LH (3 μM final) was mixed with ParC and/or MatP at a 1:2(:2) molar ratio for 30 min on ice. For experiments assaying binding of MatP, or variants, to DNA and/or LH MukB, MatP (7 μM final) was mixed with LH and/or DNA in a 1:1(:1) molar ratio and incubated on ice for 30 min. Duplicate 10 μl samples were loaded onto 10% native polyacrylamide gels poured in 125 mM Tris-HCl, pH 8.8. Following electrophoresis, gels were stained with Coomassie blue or ethidium bromide.

### Analytical Size Exclusion Chromatography (SEC)

For assessing complex formation, proteins were mixed at the indicated ratios and equilibrated in 50 mM HEPES pH 7.5, 100 mM NaCl, 1 mM MgCl_2_, 1 mM DTT and 10% (v/v) glycerol buffer for 1 h on ice. 100 μl of these mixtures (containing <900 μg of total protein) were loaded onto a Superose 6 Increase 10/300 column equilibrated in the same buffer. Separation was conducted at a flow rate of 0.5 ml/min and 0.3 ml or 0.5 ml fractions collected for SDS-PAGE analysis.

### Functional analyses *in vivo*

The ability of MukB variants to complement the temperature-sensitive growth defect of a *ΔmukB* strain was tested as described previously (13).

### Epifluorescence microscopy and Photoactivated Localization Microscopy (PALM)

The conditions for all imaging and analysis are as described in (2, 11). The analyses of singlemolecule ParC cellular diffusion and how it is impacted by the presence of MukB were as in (11). Cells for imaging were grown in M9 glycerol minimal media at 30 °C. The genotype of all strains used is described and/or cited in Figure 5.

## RESULTS

### MatP dimers and ParC dimers bind the MukB dimeric hinge domain with the same 1:1 stoichiometry

We previously established that MatP dimers bind to the dimerization ‘hinge’ domain of MukB although the precise details of this interaction were not characterised (6). Here we have used three independent assays to determine the stoichiometry and binding affinity of MatP to the MukB hinge. We exploited a MatP variant, MatPΔ18C that carries a deletion of 18 C-terminal amino acid residues because it was more amenable for biochemical studies than the full-length protein; we refer to MatPΔ18C as ‘MatP’ hereafter. Like wild type MatP, this protein is dimeric, binds *matS* sites and the MukB dimerization hinge, while a variant, MatPΔ20C, with 2 further C-terminal residues removed, retains the wild type MatP ability to displace MukBEF complexes from *ter in vivo* (6).

Initial assays used a truncated MukB variant, ‘Long Hinge’ (LH; amino acid residues 568-863), a stable dimer that encompasses the dimeric globular hinge domain and 20% of the coiled-coil region (Figure 1A; Supplementary Figure 1A-B). A 1:1 stoichiometry of binding of MatP to LH (one LH dimer binds one MatP dimer) (*K*_d_ of 0.59 μM ± 0.04) was obtained in isothermal calorimetry (ITC) assays (Figure 1B). The same 1: 1 stoichiometry was determined in native mass spectrometry (nMS) (Figure 1C) and by analytical ultracentrifugation (AUC) (Supplementary Figure 1C). To test whether MukB hinge dimerization is necessary for MatP binding, we utilised the observation that a further hinge variant, ‘Short Hinge’, lacking the entire coiled-coil (SH; amino acid residues 667-779) (Supplementary Figure 1A), forms mixtures of monomers and dimers (Supplementary Figure 1D). Complexes of MatP dimers with both SH monomers and dimers were observed in nMS (Supplementary Figure 1E), demonstrating that dimerization of the MukB hinge is not essential for MatP dimer binding.

We then analysed ParC binding to LH using ITC and nMS. In both assays, we measured a stoichiometry of 1:1 for ParC dimer-LH dimer complexes (Figure 1D-E). The affinity of the interaction, determined by ITC (0.69 ± 0.11 μM), was similar to that previously reported (7) and to the affinity between MatP and LH measured here. This result is consistent with work which showed binding of two monomeric ParC C-terminal domains to a dimeric MukB hinge comparable to LH (7). Nevertheless, our measured 1: 1 stoichiometry for complexes of ParC dimers with MukB hinge dimers contrasts with that determined in a previous study, which reported that a single MukB dimer bound two dimeric ParC molecules (7). This result was interpreted as both ParC C-terminal domains in an intact ParC dimer being unable to simultaneously bind a single dimeric hinge, because the ParC binding sites in a dimeric hinge are ~40 Å apart, whereas the ParC C-terminal domains in a ParC dimer are ~190 Å apart in the crystal structure (7, 10).

We are confident that our measurements of the 1:1 stoichiometry, using two independent assays, are robust and furthermore, nMS with a molar excess of ParC (1:8) still yielded a 1:1 stoichiometry (Supplementary Figure 1F). We have considered the possibility that in the previous stoichiometry determination using ITC, in which two ParC dimers were reported to bind a single MukB hinge dimers, not all of the ParC was active, particularly given the large variance in the enthalpies reported (7). T o accommodate the 1: 1 stoichiometry determined here, we propose that either the dimeric hinge must open and/or the C-terminal domains in a dimeric ParC must adopt a conformation in which they are closer together, which is not unreasonable given the different reported conformations of a topoIV heterotetramer (20). We conclude that MatP dimers and ParC dimers each bind to the MukB dimeric hinge domain with comparable affinities, forming complexes of 1:1 stoichiometry.

### MatP and ParC/topoIV competitively interact with the MukB hinge

Earlier *in vivo* analyses led us to infer that topoIV and MatP might compete for binding to MukB. This was because fluorescent MukBEF complexes containing the mutated variant MukB^E1407Q^ (hereafter MukB^EQ^), which binds ATP, but is hydrolysis impaired (21, 22), were enriched at *matS* sites, dependent on the presence of MatP, but not detectably associated with topoIV (4, 6, 12). In contrast, topoIV was associated with wild type ATP-hydrolysis-competent MukBEF complexes associated with the chromosomal replication origin region *(ori)* (12).

To investigate whether MatP and ParC compete for binding to the MukB hinge *in vitro,* we first tested whether MatP, ParC and LH could form ternary complexes using analytical size exclusion chromatography (SEC). Initial experiments used the monomeric ParC C-terminal domain (ParC^CTD^), which contains the MukB-binding interface (7), so that molecular mass differences between complexes would be more readily resolved. LH, MatP and ParC^CTD^ were co-incubated at a 1:2:2 ratio at μM concentrations for 1 h prior to injection. We found no evidence of ternary complexes; only the binary LH-MatP and LH-ParC^CTD^ complexes were present (Figure 2A). Similarly, no ternary (LH-MatP-ParC^CTD^) complexes were detected using nMS (Figure 2B). Consistent with these observations, when full length ParC dimers were used instead of ParC^CTD^, the binary MatP-LH and ParC-LH complexes were abundant, but only traces of possible LH-MatP-ParC complexes were detected (Figure 2C). To test whether this competition was also evident for topoIV, we reconstituted ParC_2_E_2_ heterodimers, and analysed their ability to bind to LH dimers in the presence of equal amounts of MatP. ParC_2_E_2_ complexes interacted with LH dimers, but higher-order complexes that included MatP were absent (Figure 2D). We also detected a low abundance of complexes with the stoichiometry LH4-ParC2E2; we propose these arise from higher order coiled-coil interactions, with their functional significance, if any, remaining unclear.

**Figure 2.**
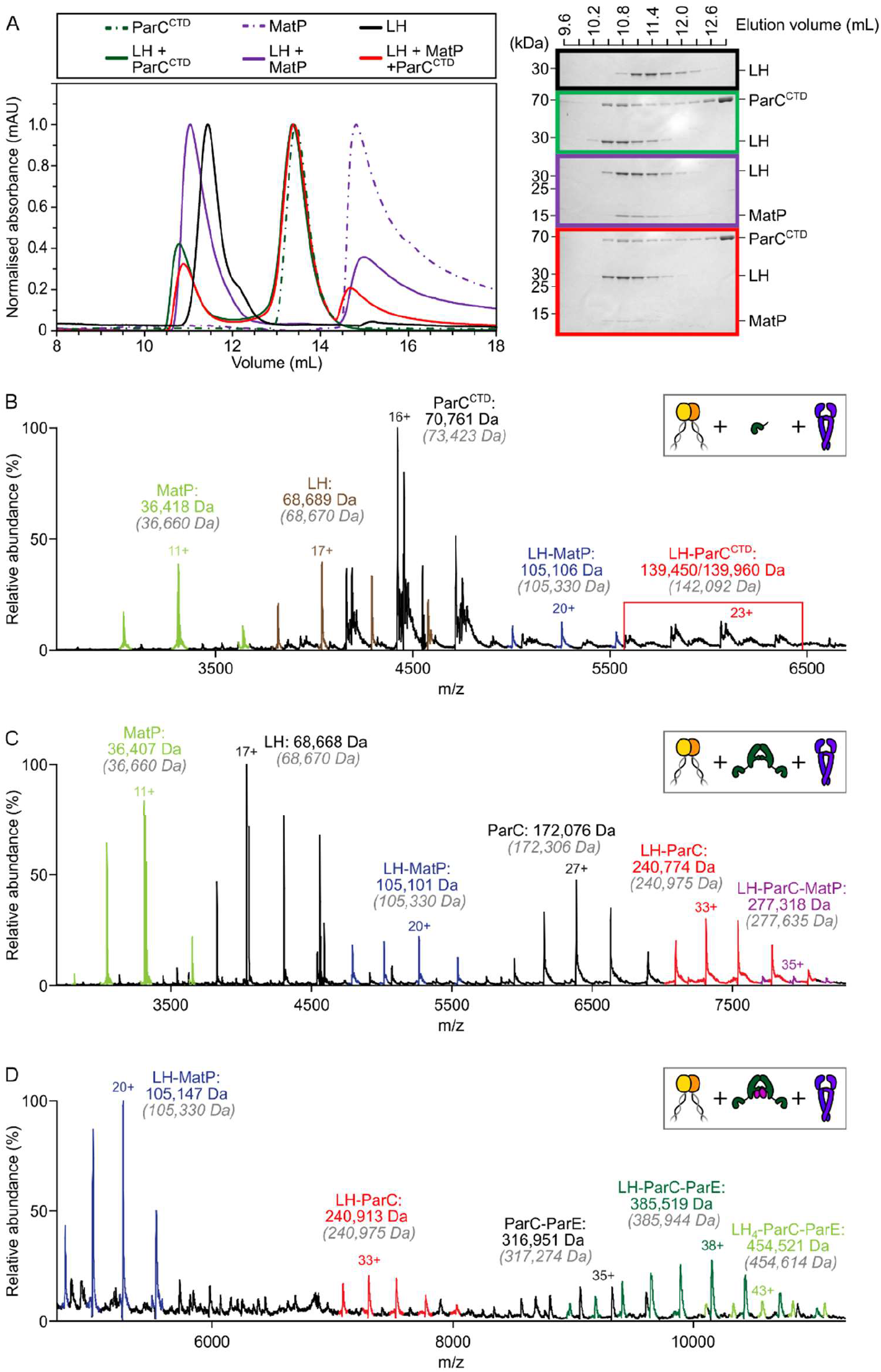
The MukB hinge does not form ternary complexes with MatP dimers and topoIV. (**A**) Analytical SEC. LH, MatP and ParC^CTD^ were co-incubated at a 1:2:2 ratio at μM concentrations for 1 h prior to injection and separated on a Superose 6 Increase column (left). 300 pL elution fractions were analysed by SDS-PAGE and Coomassie staining (right). Note ParC^CTD^ retains its MBP-N-terminal fusion (~46 kDa) for reasons of stability (see Materials and Methods). (**B-D**) Representative mass spectra of complexes detected when co-incubating LH and MatP in 1:2 ratio with 2 equivalents of either ParC^CTD^ (**B**), ParC alone (**C**) or topoIV heterotetramers (**D**) for 10 min on ice. Grey italics denote the theoretical mass of complexes.

To analyse the competition between ParC and MatP for binding to the MukB hinge more quantitatively, we exploited FCS, using Cy3B-labelled MatP at a cysteine residue added to the His-tag at the C-terminus (MatP has no intrinsic cysteines). MukB bound MatP-Cy3B with a similar affinity (*K*_d_ ~0.25 μM) to that observed for LH binding to MatP (Figure 3A, Supplementary Figure 2A). This indicates that the conformation of the hinge in LH is comparable to that of the hinge in full-length MukB. As expected, the interaction between MatP-Cy3B and MukB was competed out when a 50-fold molar excess of unlabelled MatP was added. A 3-fold decrease in binding was observed when the reaction was challenged with a 10-fold excess of either ParC or ParC^CTD^ (Figure 3A), while ParC^R705E^,^R729A^ (ParC^EA^), a mutant defective in MukB hinge binding (8) (Supplementary Figure 2C), did not significantly decrease MatP-Cy3B binding to MukB. No interactions between MatP and ParC were detected by FCS or nMS (Supplementary Figure 2D-E).

**Figure 3.**
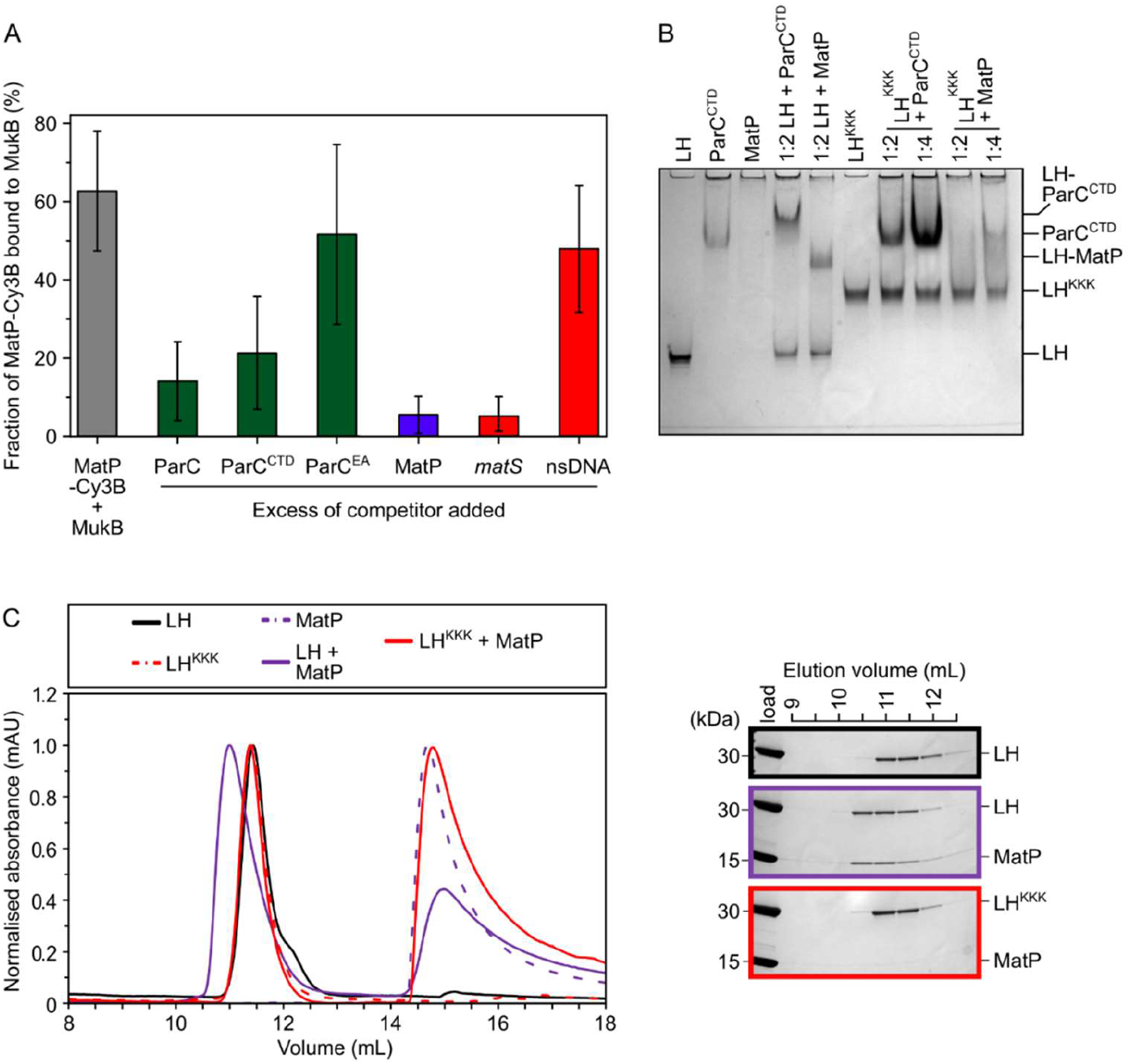
MatP and ParC compete for binding to overlapping sites on the MukB hinge. (A) FCS measurements of competition for binding between ParC and MatP for MukB and also between MatP and 15 bp DNA hairpins, containing a 13 bp *matS2* site or non-specific sequence, for MukB. Cy3B-labelled [MatP] was fixed at 1 nM and wild type [MukB] at 400 nM; this achieved ~60% binding of all MatP-Cy3B to MukB, consistent with their measured *Kd* (Supplementary Figure 2A). The diffusion times of free MatP-Cy3B, MukB-TAMRA and MukB-bound MatP-Cy3B were determined to be ~4.5 ms, ~15 ms and ~ 20-50 ms, respectively, allowing identification of the MukB-MatP population. Auto-correlation curves were fit to a two-component equation (Equation 2) with the diffusion time of Cy3B-MatP fixed to 4.5 ms (*τ*_1_), whilst the diffusion time (*τ*_2_) of bound complex was allowed to float to obtain the best fitting for the data. All ParC variants, unlabelled MatP and DNA were added at a 10-, 50-, and 2.5-fold molar excess, respectively. Error bars represent (mean ±SD). (B) Native PAGE. Varying ratios of MatP or ParC^CTD^ were incubated with LH (at 3 μM) for 30 min on ice prior to electrophoresis under non-denaturing conditions. All proteins were run alone as a reference for their mobility in an 8% tris-glycine gel. Note, MatP alone poorly enters the gel. (**C**) Analytical SEC. Either wild type LH or the LH^KKK^ variant (already characterised to be defective in ParC binding) were co-incubated with MatP at a 1:2 ratio with LH (at 20 μM) for 1 hr on ice prior to injection and separated on a Superose 6 Increase column (left). 500 pL elution fractions were analysed by SDS-PAGE and Coomassie staining (right).

Given the demonstration that ParC and MatP compete for binding to the MukB hinge, we explored whether MatP and ParC share the same or overlapping binding sites on the hinge, by analysing variants of LH and full length MukB that were known to be deficient in ParC binding (9). LH and MukB derivatives containing a triple substitution (D697K D745K E753K; LH^KKK^ and MukB^KKK^ thereafter) were impaired in both ParC and MatP binding when the interactions were assayed by native PAGE (Figure 3B), SEC (Figure 3C) and FCS (Supplementary Figure 2A), respectively. In the latter case the *K*_d_ was increased from ~0.25 μM to ~1.6 μM. This defect was not attributable to global misfolding (Supplementary Figure 2B). We conclude that topoIV and MatP compete for binding to the MukB hinge as a consequence of MatP and ParC having overlapping binding sites on the hinge.

### The MukB hinge fails to stably bind MatP-matS complexes

Since the action of MatP in displacing MukBEF complexes from *ter in vivo* requires that MatP is at least transiently bound to *matS* sites (2, 6), we initially anticipated that MatP-matS complexes would form stable complexes with the MukB dimerization hinge. We were therefore surprised to see that addition of an excess of a 50 bp DNA fragment carrying an internal 13 bp *matS2* (23) site almost totally abolished binding of MatP-Cy3B to MukB in FCS, while a non-specific DNA fragment of the same length and GC content had no detectable effect on binding (Figure 3A). Furthermore, when we incubated MatP-matS complexes with LH, we failed to observe ternary LH-MatP-matS complexes in nMS, although the binary MatP-matS and LH-MatP complexes were present (Figure 4A). We confirmed that under these conditions, MatP was specifically bound to *matS* (Supplementary Figure 3A). Addition of topoIV failed to stabilise complexes of MatP-matS with LH (Supplementary Figure 3B). Similarly, in native PAGE, we identified the binary, but not the ternary complexes (Figure 4B). Again, the presence of the DNA fragment that did not bind MatP specifically, did not fully impair MatP binding to LH (Figure 4B, Supplementary Figure 3C); note that non-specific DNA binding to MatP during electrophoresis led to smeared complexes containing both MatP and DNA. We detected no stable complexes of LH with 50 bp DNA in nMS (Supplementary Figure 3D).

**Figure 4.**
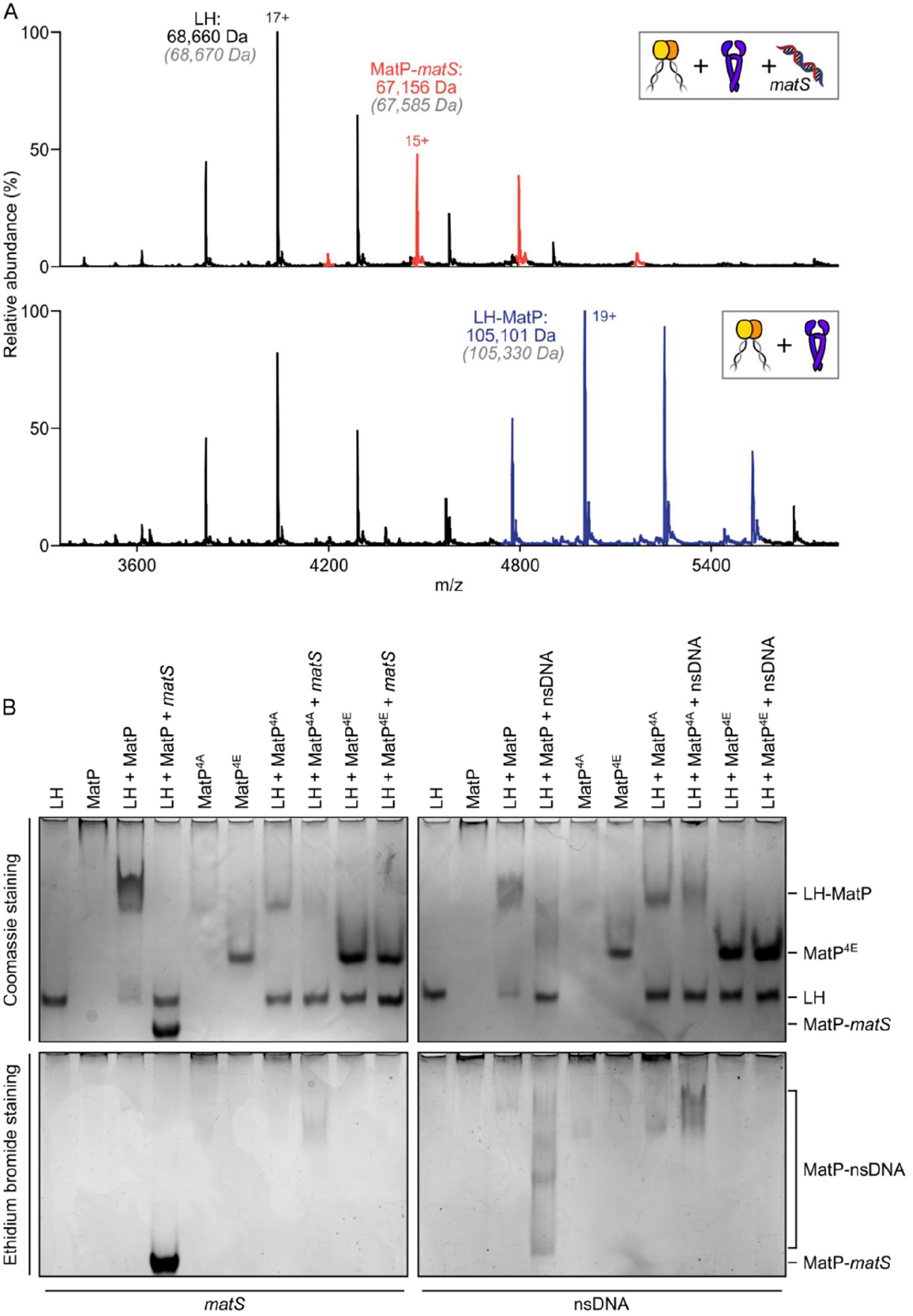
*matS* sites compete with the hinge for MatP binding. (**A**) Representative mass spectra of species detected between LH and MatP in the presence of *matS* DNA. LH, MatP, and in the case of the upper panel *matS,* were mixed at 1:2:1. Grey italics denote the theoretical mass of complexes. (**B**) Native PAGE. Formation of LH-MatP-matS ternary complexes were not detected. MatP^4A^ (K71A, Q72A, R75A and R77A) and MatP^4E^ (K71E, Q72A, R75E and R77E) are impaired in binding to the MukB hinge domain. The same samples were loaded onto two equivalent native gels – one for Coomassie staining and one for ethidium bromide staining.

Since small angle X-ray scattering (SAXS) analysis of the MatP envelope indicated that MatP alone is less compact than MatP-matS, whose structure has been determined by X-ray crystallography (24), we considered whether these global conformational differences are responsible for the observed differential binding to the hinge. However, given that MatP binding to *matS* and LH is mutually exclusive, we explored the alternative hypothesis that *matS* and the MukB hinge share an overlapping binding interface on MatP. The structure of MatP-matS identified residues involved in DNA binding, including K71, Q72, R75, and R77 (24). Consistent with this, a quadruple substitution of these residues with Ala (MatP^4A^) which neutralised their charge, or with Glu (MatP^4E^) which reversed it, led to a loss of specific binding with *matS* DNA (Supplementary Figure 3C); though MatP^4A^ retained partial ability to interact with a non-specific DNA fragment (Supplementary Figure 3C). Both of these MatP variants were impaired in their interaction with LH, consistent with MatP using overlapping determinants to bind MukB hinge and *matS* (Figure 4B). It is also possible that MatP^4A^ and MatP^4E^ might have altered overall structures that prevent hinge binding; nevertheless, they are both dimeric and generated a Gaussian distribution of charge states in nMS, indicative of folding (Supplementary Figure 3E). Therefore, we favour a model in which *matS* and the hinge share a common binding interface on MatP.

### Cells expressing MukB hinge mutants defective in binding of ParC and MatP *in vitro* are Muk^+^ and impaired in ParC and MatP interactions *in vivo*

Given that the MukB^KKK^ hinge mutant analysed above is deficient in both ParC and MatP binding *in vitro,* we anticipated that cells expressing MukB^KKK^ might exhibit both a MatP’ phenotype and a chromosome segregation-cell division phenotype, resulting from the lack of correct targeting and catalysis by ParC/topoIV. Moreover, we considered that this mutant might additionally have a MukB’ temperature-sensitive growth phenotype, since temperature-sensitive growth was reported for a different MukB hinge mutant (MukB^D692A^) that was also impaired in ParC binding (7).

*ΔmukB* cells with fluorescently labelled *ori1* and *ter3* loci and a functional chromosomal *mukE-mYPet* gene (2, 6), expressing basal levels of wild type MukB from multicopy plasmid pBAD24, formed ori-associated fluorescent MukBEF foci when grown at 30 °C in minimal glycerol medium (Figure 5A-B), as expected, given that they were not temperature-sensitive (Figure 5C). Negative control *ΔmukB* cells containing just the pBAD24 vector, showed no evidence of nucleoid-localised fluorescent MukBEF complexes, consistent with their temperature-sensitive growth. *ΔmatP* cells exhibited distinct MukBEF foci that were positioned equally distant from *ori1* and *ter3,* as reported previously, because of the failure to deplete MukBEF from *ter* (2, 6). The preferential ori-association of MukBEF complexes in wild type cells arises directly from their depletion from *ter* as a consequence of the ATP hydrolysis-dependent MukBEF dissociation from the 23 MatP-bound *matS* sites within *ter* (2). Cells expressing basal levels of MukB^KKK^ in the *ΔmukB* chromosomal background, were not temperature-sensitive (Figure 5C) and formed somewhat diffuse fluorescent MukBEF complexes (Figure 5A), which showed a more distant localization with the *ori1* locus (with similar distances between the MukBEF complexes and *ter3)* (Figure 5D). This phenotype, in which preferential *ori1* localisation was lost, is similar to that of *ΔmatP* cells (Figure 5D) (2) and is consistent with reduced interaction of MukB^KKK^ with MatP *in vivo.* Nevertheless, the phenotypes of MukB^KKK^ and *ΔmatP* cells were not identical, with the MukBEF foci in the latter cells being discrete, while those in the MukB^KKK^ cells being somewhat diffuse, though unlike the uniform distribution of molecules in Muk’ cells (Figure 5A). Note that in *ΔmukB* cells, the image analysis software identifies the brightest MukE pixel in every cell and measures the distance from it to the *ori1* and *ter3* loci. Since we expect this pixel to be placed randomly in the cell, given the uniformly distributed fluorescence (Figure 5A), the mean measured distances to *ori1* and *ter3* are expected to be identical, as was observed (Figure 5D). We conclude that MukB^KKK^EF complexes are functional given that cells expressing these are not temperature-sensitive and form chromosome-associated complexes. Nevertheless, these no longer preferentially associate with *ori1* as a consequence of a reduced association with MatP-matS at *ter.*

**Figure 5.**
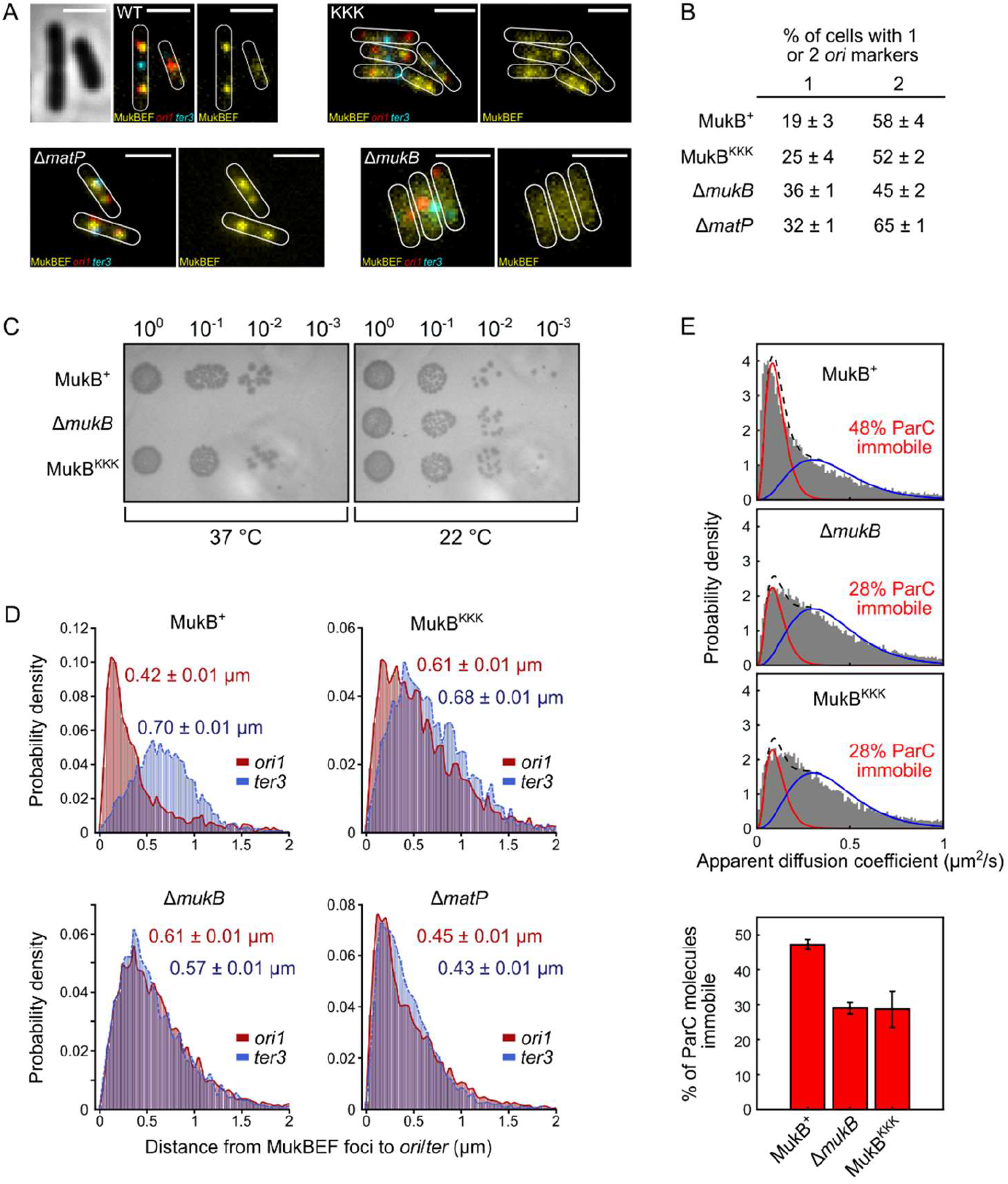
Cells expressing MukB^KKK^ are impaired in ParC and MatP binding. (**A**) Representative fluorescence images with cell borders of *ΔmukB* cells with fluorescently labelled MukE (mYPet), ori1(mCherry), and *ter3* (mCerulean) (AU2118; *lacO240 @ori1* (3908) *(hyg), tetO240@ter3* (1644) (gen), *ΔleuB::Plac-lacI-mCherry-frt, ΔgalK::Plac-tetR-mCerulean-frt, mukE-* mYPet ΔaraBAD::FRT (AraC+) FRT-T1-T2-Para-ΔmukB::kan) (2, 6), expressing basal levels of pBAD24 plasmid-borne WT MukB, MukB^KKK^, and empty pBAD24 plasmid control *(ΔmukB). ΔmatP* cells expressing MukBEF under the native *smtA-mukBEF* promoter, with fluorescently labelled MukE, *ori1,* and *ter3* labelled MukB, *ori1* and *ter3* (SN302) (2) Scale bars: 2 pm. (**B**) Percentage of cells in (**A**) with either 1 or 2 *ori1* markers (± SEM). (**C**) MukBEF phenotype of MukB^+^, MukB^KKK^ and *ΔmukB* cells as judged by temperature-sensitive growth in rich medium (LB) at 22 °C and 37 °C. Basal levels of plasmid-borne MukB, MukB^KKK^ were expressed from cells carrying a chromosomal MukB deletion (PZ129) (11); *ΔmukB* control cells carried the empty plasmid. (**D**) Distances between MukBEF foci (measured by the brightest pixel; 2,6) and *ori1/ter3* markers (±SEM) in the strains in (A). WT MukB, 4837 cells, median cell length 2.87 μm; MukB^KKK^, 5846 cells, median cell length 2.98 μm; *ΔmukB,* 10,670 cells, median cell length 2.76 μm, *ΔmatP,* 17,933 cells, median cell length 3.10 μm. (**E**) Singlemolecule tracking (PALM) of ParC-PAmCherry molecules in *ΔmukB* cells (PZ129) (11) complemented with basal levels of plasmid-expressed WT MukB or MukB^KKK^; control *ΔmukB* cells contained an empty plasmid. For each condition, the distribution of ParC apparent diffusion coefficients was fitted to a 2-species model as in (11). Bar chart shows same data, with SD from 3 experimental repeats.

Furthermore, MukB^KKK^ expressing cells showed no evidence of defects in decatenation/chromosome segregation resulting from impaired action of topoIV, since they had almost identical cell size distributions to those expressing WT MukBEF and an *ori1* focus number distribution more similar to WT and *ΔmatP* cells, than to *ΔmukB* cells, which have a well characterized delay in segregation of newly replicated *ori1* loci (Figure 5D) (4, 6). Interaction of topoIV molecules with immobile chromosome-associated MukBEF complexes *in vivo,* leads to a higher proportion of topoIV molecules becoming immobile (11). Using an identical analysis, we compared the ParC singlemolecule distribution in MukB^KKK^, MukB^+^ and MukB’ cells using PALM (Figure 5E). Cells expressing MukB^KKK^ showed an almost identical distribution of immobile/mobile ParC molecules to cells lacking MukBEF, thereby demonstrating that the topoIV-MukB interaction is ablated *in vivo* in the Muk^KKK^ mutant. We conclude that the failure of ParC to bind MukB *in vivo* does not seriously impact the ability of cells to mediate topoIV-mediated decatenation.

## DISCUSSION

The results presented here extend our understanding of the important functional interplay between MukBEF, MatP and topoIV in the organising and processing of *E. coli* chromosomes. Previous work has shown that ParC dimers and topoIV ParC_2_E_2_ heterotetramers bind to the MukB dimerization hinge through one of the five ParC C-terminal ‘blades’, the interaction leading to stimulation of topoIV catalysis (7, 9, 10). Interaction of this same C-terminal blade with transfer segment DNA, may direct the capture of a specific DNA topology during strand transfer, with MukB hinge binding leading to impairment of the interaction between ParC^CTD^ and DNA *in vitro* (10). Our demonstration that binding of MatP dimers and ParC dimers/topoIV heterotetramers to the dimeric MukB hinge is mutually exclusive provides an attractive potential means for the spatial and temporal regulation of topoIV activity by MukBEF and MatP. Consistent with this competition is our observation that mutations in the MukB hinge that ablate ParC binding also severely impair MatP binding *in vitro,* and with the *in vivo* observation that topoIV was not associated with MukB^EQ^ complexes enriched at MatP-matS sites within *ter* (12).

How can we reconcile the demonstration here, using two independent assays, that only one intact ParC dimer, or a topoIV heterotetramer binds a single dimeric hinge, when earlier work indicated that two ParC dimers bind one hinge dimer (7)? To explain the earlier result, it was proposed that a steric constraint prevents the two C-terminal domains of ParC, ~190 Å apart in the ParC crystal structure, from docking on the two binding sites separated by ~45 Å on a dimeric hinge, thereby leading to two ParC dimers bound to the hinge, with each having only one of its two CTDs bound (10). Nevertheless, even with an 8-fold molar excess of ParC in our nMS assay, we still observed only 1:1 ParC dimer: MukB hinge dimer complexes, with no evidence of 2:1 complexes, strengthening the conclusion that 1:1 is the physiologically relevant stoichiometry. The most logical explanation of this is that a single ParC dimer, either alone, or in a topoIV heterotetramer, has both of its C-terminal domains interacting with the two binding interfaces on a single dimeric MukB hinge. An alternative explanation, in which binding of one ParC^CTD^ of a ParC dimer to one side of the hinge is incompatible with binding of a second ParC^CTD^ to the second hinge interface, because of negative cooperativity or steric constraints, seems unlikely given that two isolated ParC CTDs bind independently to the two hinge binding interfaces (7, 10). Our favoured explanation requires that the ParC C-terminal domains adopt an alternative conformation in which they are closer together and/or the binding interfaces on the MukB hinge move apart by breaking or reorganising the dimerization interface. Precedents for alternative conformations of the ParC C-terminal domains (and their equivalent in other topoisomerases) relative to the core enzyme have come from structural and modelling analyses of other bacterial type II topoisomerases (20, 25–28). Furthermore, hinge opening on association with a ParC dimer would be consistent with previous proposals for other SMC complexes suggesting hinge opening to allow DNA passage into or from the SMC ring (29–31). Assuming our proposal is correct, it leads to a scenario in which a topoIV heterotetramer bound to a MukB hinge will have one of its DNA passage gates sitting positioned above the hinge, whose own gate, formed by dimerization, potentially opening to allow DNA passage between topoIV and MukBEF. It is attractive to think that in these two ‘multi-gate’ protein complexes, regulated gate opening and closing may be used for coordinating topoIV and MukBEF action.

We were initially surprised to observe that matS-bound MatP failed to bind stably to the MukB hinge, because we previously established that MukBEF complexes detect MatP bound to *matS* sites *in vivo,* leading to MukBEF dissociation and depletion from *ter* (1, 2, 6). Our demonstration that MatP, but not MatP-matS, binds the MukB hinge stably and that MukB hinge and *matS* DNA compete for binding to MatP may reflect a transient ‘handover’ state related to MukBEF dissociation from the chromosome. A comparable ‘handover’ state may explain the observation that DNA and the MukB hinge compete for binding to ParC *in vitro,* despite their *in vivo* action presumably requiring the participation of all three components. Indeed, it seems that the acidic hinge binding interface for MatP and ParC is effectively a DNA mimic. As is often the case, our *in vitro* biochemical assays reflect a ‘snapshot’ rather than complete *in vivo* behaviour, which in the case of MukBEF involves very large multiprotein dimer of dimer complexes, whose conformations may well involve the MukB hinge associating with the ATPase heads, through bending of the coiled coils at an elbow (32). Nevertheless, we are confident that the interaction of MatP with the MukB hinge characterized here is functionally relevant, given the specificity and affinity of binding with defined stoichiometry in different assays, and the inferred impaired association of the MukB^KKK^ variant with MatP-matS *in vivo.*

The work here highlights the importance of combining different biochemical assays with quantitative *in vivo* analysis. Our demonstration that cells expressing the MukB^KKK^ variant fail to associate with topoIV *in vivo* is reassuring and validates the results from *in vitro* assays reported here and elsewhere (7–10). Nevertheless, we were surprised that cells expressing this variant showed no obvious topoIV-defective phenotype as assessed by normal cell division and no observable defect in segregation of newly replicated oris. We have proposed previously (2) that some or much of the Muk’ phenotype, including the delayed segregation of newly replicated oris, arises from defective decatenation. The data here do not support that proposal. The substantial body of work that has characterized the MukB-topoIV interaction can be compared with the many proposals elsewhere that implicate a range of SMC complexes in acting together with type II topoisomerases (33–37), although in the latter cases there is no direct evidence to identify the type of interaction. Although bacterial topoIV is the major decatenase, the type I topoisomerase, topoIII, can decatenate regions of chromosomes containing single strands, for example at replication forks (38, 39). Furthermore, FtsK-dependent XerCD site-specific recombination at *E. coli dif* locus can efficiently remove replicative catenanes within *ter* (40, 41), while loss of *E. coli* FtsK translocation activity combined with a lack of functional MukBEF leads to extensive filamentation, chromosome segregation defects and inviability (42), possibly because of a combined decatenation defect in such cells. Intriguingly, *Bacillus subtilis* SMC is not known to interact with topoIV, but rather it does interact with the recombinase XerD, to facilitate recombination-independent expulsion of SMC complexes from *ter* (43). Additionally, it is conceivable that the *B. subtilis* SMC-XerD interaction promotes decatenation by XerCD recombination at *dif* in that organism. The demonstration that *B. subtilis* SMC complexes are displaced from *ter* in this organism by interaction with ter-bound XerD (43) underlines the fact that displacement of these complexes from *ter* is not restricted to the few bacterial species that encode MukBEF-MatP and that such displacement may be a general feature of bacterial chromosome dynamics.

## DATA AVAILABILITY

All digital forms of the data are available on request. All materials and analysis codes are available upon reasonable request.

## SUPPLEMENTARY DATA

Supplementary Data are available at NAR online.

## AUTHOR CONTRIBUTIONS

L.K.A., G.L.M.F., K.V.R. and D.J.S. conceived and directed the project. J.R.B., R.B., G.L.M.F., K.V.R. and M.Z. undertook biochemical experiments. J. M., J.P.P. and M.S. undertook quantitative imaging and analysis. C.V.R. provided facilities for nMS. The paper was drafted by L.K.A., G.L.M.F. and D.J.S., with all authors participating in the final manuscript.

## ACKNOWLEDGEMENT

We thank all members of the Sherratt lab for useful discussions and David Staunton for expert technical assistance with ITC. Achillefs Kapanidis (Dept of Physics, University of Oxford) provided facilities for FCS analysis. We thank Madhu Srinivasan (Biochemistry Oxford), Frank Burmann and Jan Lowe (MRC LMB, Cambridge, UK) for helpful discussions. We also thank K. Zawadzka and P. Zawadzki (MNM, Poznan, Poland), whose actions and ideas helped initiate the work in this paper.

## FUNDING

This work was supported by a Wellcome Investigator Award [200782/Z/16/Z to D.J.S.; 104633/Z/14/Z]. An MRC Programme Grant [MR/N020413/1] awarded to C.V.R. supported the native mass spectrometry. Funding for open access charge: Wellcome Trust [200782/Z/16/Z].

## CONFLICT OF INTEREST

None declared.

---

**Supplementary Figure 1.**
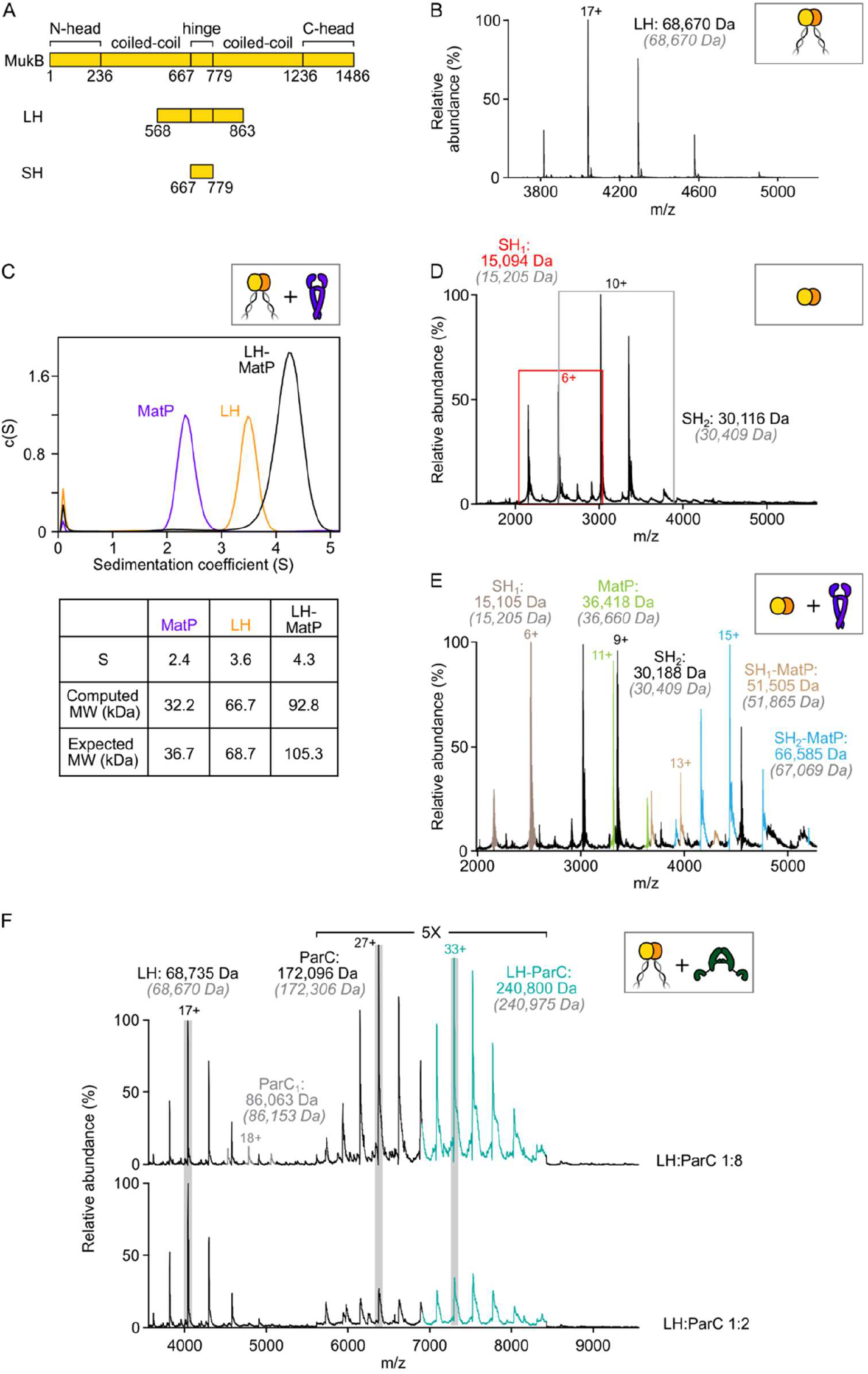
MatP dimers interact with the MukB hinge domain. (**A**) Schematic of the functional domains of MukB and hinge-based truncations (long hinge (LH) and short hinge (SH)). (**B**) nMS confirming that LH is a stable dimer. (**C**) Sedimentation velocity (SV-AUC) experiments showing complex formation between MatP and LH. (**D**) Mass spectrum showing SH MukB is a monomer/dimer mix at tens of μM concentrations. Peaks corresponding to monomeric species are boxed in red. Grey italics denote the theoretical mass of complexes. (**E**) Representative spectrum showing binding of MatP to SH MukB. (F) nMS titration experiment with ParC at a 2-fold and 8-fold molar excess relative to LH. In both cases, a 1:1 LH:ParC complex is detected only.

**Supplementary Table 1.**
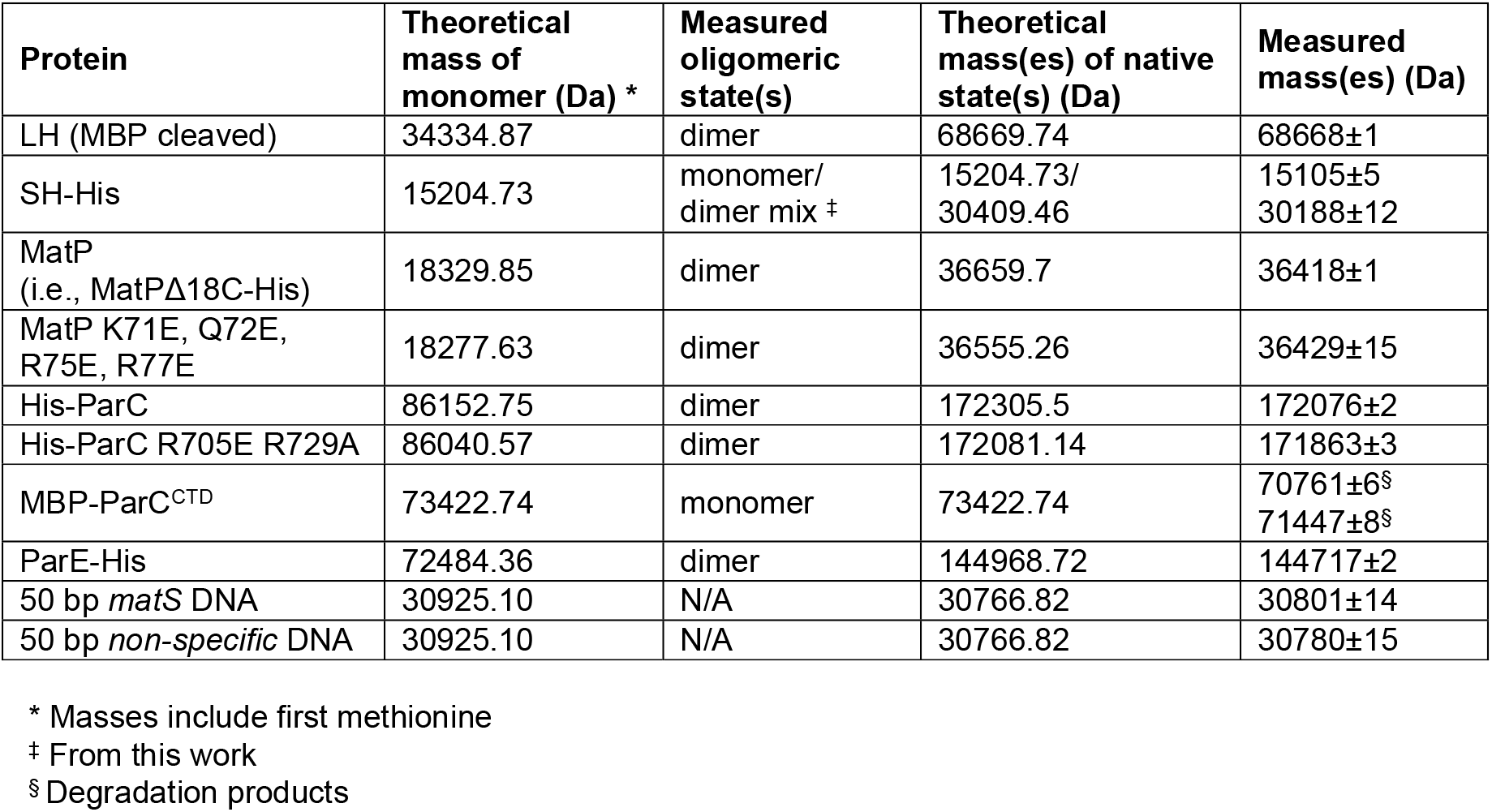
Theoretical masses of proteins and DNA used in native MS. Predicted and measured masses of MukBEF, topoIV and MatP components or variants and also DNA substrates. Errors are the standard deviation in mass determination.

**Supplementary Figure 2.**
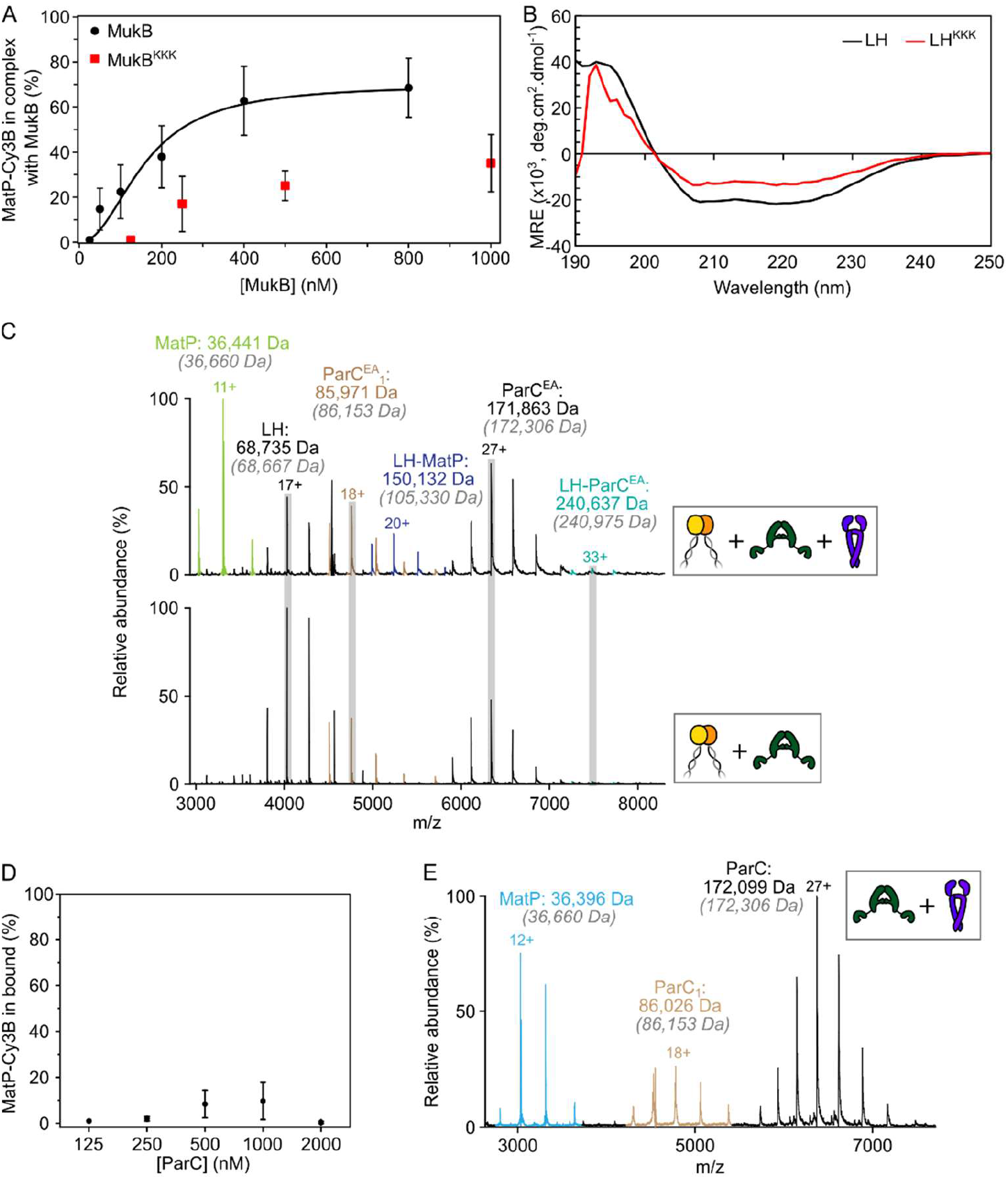
MatP and ParC compete for binding to the same or overlapping sites on the MukB hinge. (**A**) Determination of binding affinity of full-length MukB and MukB^KKK^ using Cy3B-labelled MatP in FCS. Titrations were performed at a constant concentration of MatP-Cy3B (1 nM). Complex percentage was set as 99% at 25 nM MukB for optimal fitting. The data were fitted to the Hill equation. Error bars (mean ± SD). (**B**) CD spectra of wild type LH and the LH^KKK^ variant. (**C**) Representative mass spectra confirming that the previously characterised ParC^R705E^,^R729A^ (ParC^EA^) variant (8) is defective in binding to the MukB hinge domain. **(D**) FCS titration assay. ParC does not interact with MatP directly. (**E**) Representative nMS spectrum demonstrating that ParC and MatP do not physically interact. Grey italics denote the theoretical mass of complexes.

**Supplementary Figure 3.**
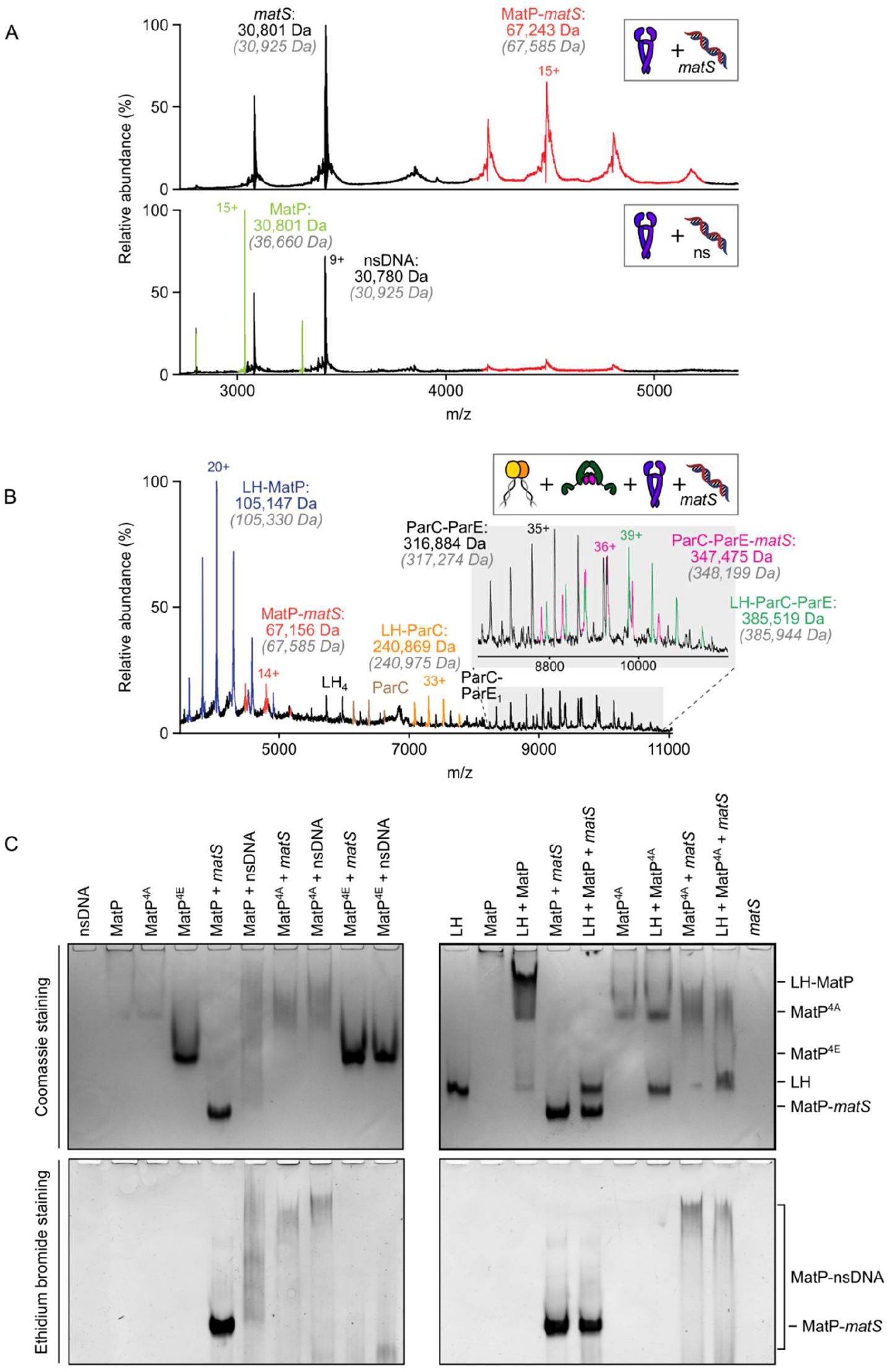
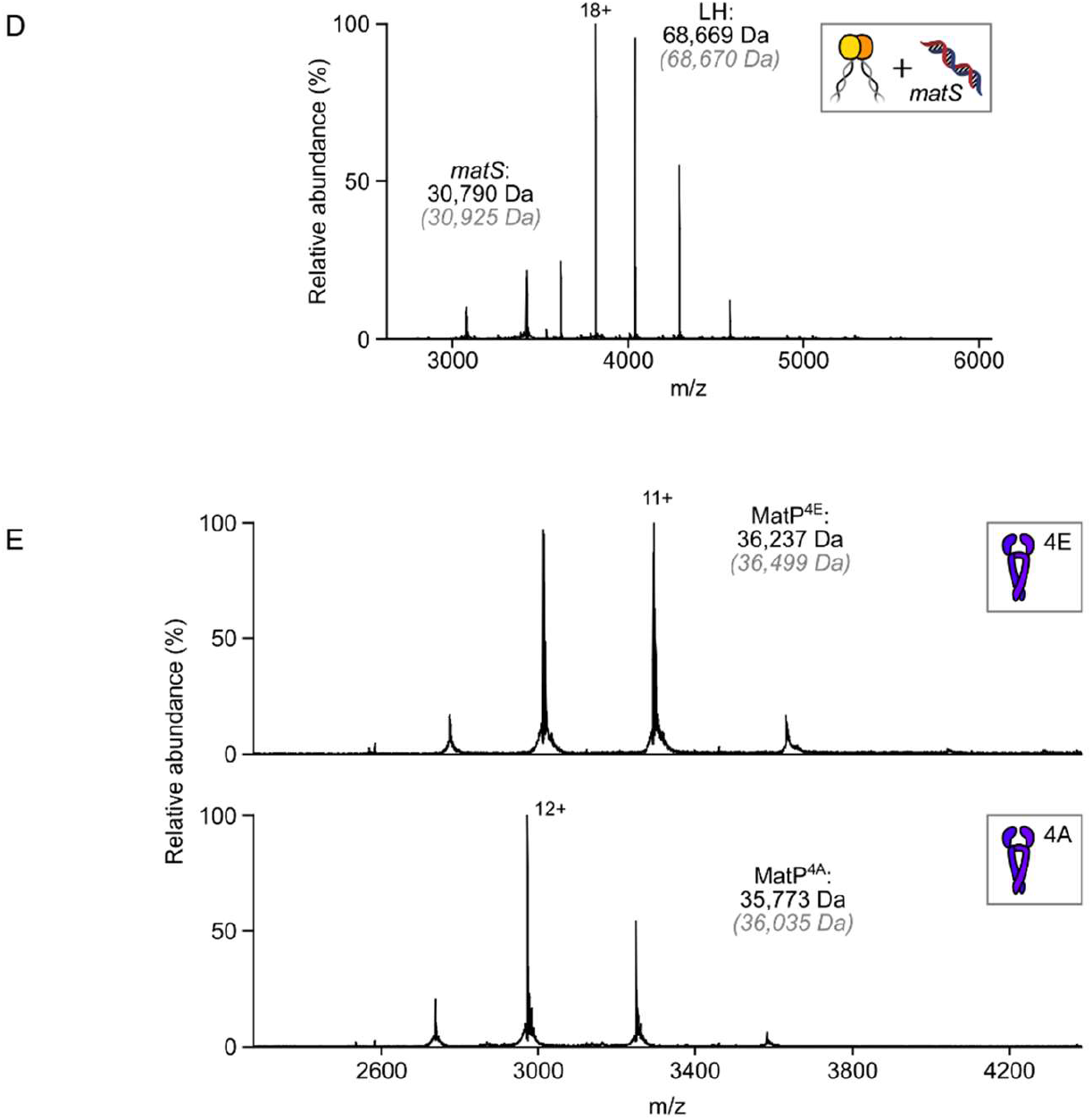
*matS* sites compete with the MukB hinge for MatP binding. (**A**) Representative mass spectra indicating that 50 bp matS-containing DNA specifically binds MatP dimers under the conditions used. MatP and DNA was mixed at 1:1. Grey italics denote the theoretical mass of complexes. (**B**) Representative mass spectra demonstrating that MatP when bound to *matS* does not stably interact with either the hinge domain (LH) of MukB or LH-topoIV complexes. (**C**) Native PAGE analysis of impaired LH and matS/non-specific DNA binding by MatP^4A^ (K71A, Q72A, R75A and R77A) and MatP^4E^ (K71E, Q72E, R75E and R77E). Gels were stained with Coomassie blue and ethidium bromide to identify nucleoprotein complexes. (**D**) nMS experiments ascertained that the hinge domain of MukB cannot stably bind 50 bp DNA. (**E**) nMS confirms that MatP^4A^ and MatP^4E^ variants are stable dimers. Additionally, their Gaussian distribution of charge states is indicative of the folded nature of these variants.

## Notes

### Competing Interest Statement

The authors have declared no competing interest.

